# 3DGV: Immersive Exploration of 3D Genome Structures using Virtual Reality

**DOI:** 10.1101/855379

**Authors:** Éric Zhang, Chrisostomos Drogaris, Antoine Gédon, Aaron Sossin, Rajae Faraj, Haifen Chen, Yan Cyr, Jacek Majewski, Mathieu Blanchette, Jérôme Waldispühl

## Abstract

The analysis of 3D genomic data is expected to revolutionize our understanding of genome organization and regulatory mechanisms. Yet, the complex spatial organization of this information can be difficult to interpret with 2D viewers. Virtual Reality (VR) technologies offer an opportunity to rethink our methods to visualize and navigate 3D objects. In this paper, we introduce the Virtual Reality 3D Genome Viewer (`3DGV`), an open platform to experiment and develop VR solutions to explore 3D genome structures.

**Availability:** http://3dgv.cs.mcgill.ca/

## Introduction

As chromosome conformation capture (3C) technologies continue to improve, the need for applications and software to process and review 3D genomic data is growing (1). In particular, the release of the first complete 3D models (2) suggests the development of customized tools to exploit this novel information.

Currently, software designed to explore 3D genome structures is primarily desktop-based and distributed as stand-alone applications (3–5) or integrated in a Web browser (6–8). But the browsing experience is limited by the representation of 3D objects as 2D projections on a screen and the manipulation of the data through keyboards and mice. Immersive technologies present an opportunity to address this challenge (9, 10).

In this paper, we introduce the Virtual Reality 3D Genome Viewer (`3DGV`) – a virtual reality (VR) version of our previous 3D genome browser (6). Notably, `3DGV` introduces multiple concepts and techniques to facilitate the manipulation of the data in immersive environment, as well as visual representations allowing to overlay multiple levels of genomic annotations in 3D space. This software aims to lower the barrier to entry to immersive technologies for 3D genomics and serve as a base for further experimentations.

## Methods

### Input data

The Virtual Reality 3D Genome Viewer (`3DGV`) displays *pre-computed* 3D models of genome structures. In addition, users can also upload genomic annotations they wish to analyze in this structural context. Sequence data is stored in a basic text file. For structure coordinates and other annotations, `3DGV` uses a simple tab-separated values (TSV) file format that enables us to easily convert annotations from any file format. We represent 3D models as a trajectory where each row of the file contains a node identifier, its genome index and spatial coordinates. A gene annotation file stores in each row the chromosome number, start and end indices, name and orientation of a gene. Other annotations use a generic format in which each row store a chromosome number, start and end indices, label and value associated to this region. The current distribution features 3D models of the 23 Human chromosomes from (11) with gene annotations stored in separate files. We also include CpG islands and Methylation data to illustrate the versatility of the file format and test `3DGV` functionalities.

### Visualization and annotations

3D structures of chromosomes are presented as 3D splines on the top of which we overlay annotations (See Fig. 1a). Genes possess their own level of representation. We display the genomic region they occupy with a cylinder wrapped around the spline, where the color gradient show the orientation of this gene. Other annotations are represented at a third level with larger cylinders whose color encode the value associated to this segment. For instance, the relative amount of methylation of regions of a chromosome. Multiple annotations can be combined and mapped on the same 3D model, but overlaps can occur at the third level. We do not display the sequence data.

**Fig. 1.**
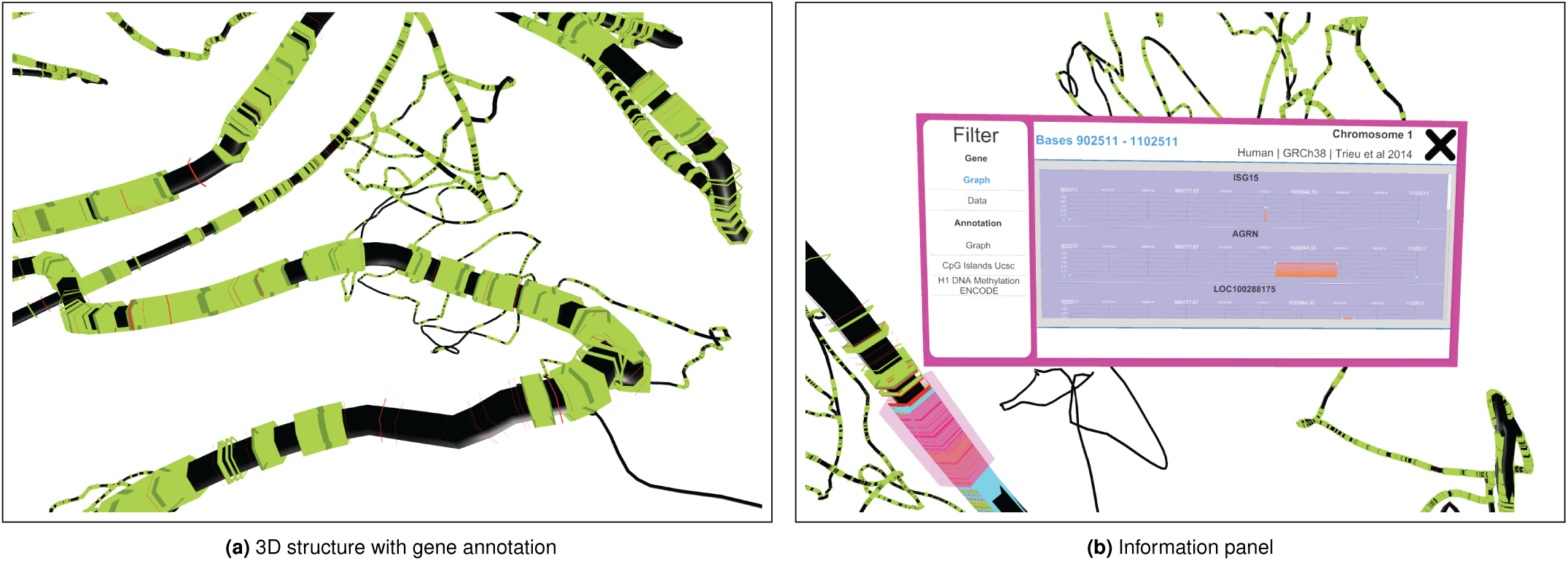
(Left) 3D structures with annotated with gene (in green). The darker green ring shows the start of the gene. (Right) Information panel showing gene positions, CpG and Methylation data of the region highlighted in Magenta.

### Navigation and Controls

By default, users navigate the 3D space and manipulate data with two controllers. The main menu allows the users to activate the 3D models and annotations to display. The speed of motion and resolution of the model (i.e zoom) can be adjusted. We also implemented a search function allowing the user to access immediately specific coordinates. During the exploration of the data, a pointer enables the users to query specific regions of the genome and display the information associated within a panel (See Fig. 1b). These panels feature a graphical summary and can be conveniently organized to compare annotations in distinct regions.

In order to improve accessibility, keyboard and mouse functionality was added to the application in order to allow users who do not have access to virtual reality to use `3DGV`.

### Implementation and Distribution

The Virtual Reality Genome Viewer (`3DGV`) is distributed as a standalone application built using the Unity game engine that works with most popular commercial VR headsets and controllers. The current release has been tested on HTC Vive and Oculus Go. It can also be used on desktops using a non-VR mode under OSX and Windows operating systems.

